# Cell-type-specific propagation of visual flicker

**DOI:** 10.1101/2023.01.04.522738

**Authors:** Marius Schneider, Athanasia Tzanou, Cem Uran, Martin Vinck

## Abstract

Rhythmic flicker stimulation has gained interest as a treatment for neurodegenerative diseases and a method for frequency tagging neural activity in human EEG/MEG recordings. Yet, little is known about the way in which flicker-induced synchronization propagates across cortical levels and impacts different cell types. Here, we used Neuropixels to simultaneously record from LGN, V1, and CA1 while presenting visual flicker stimuli at different frequencies. LGN neurons showed strong phase locking up to 40Hz, whereas phase locking was substantially weaker in V1 units and absent in CA1 units. Laminar analyses revealed an attenuation of phase locking at 40Hz for each processing stage, with substantially weaker phase locking in the superficial layers of V1. Gamma-rhythmic flicker predominantly entrained fast-spiking interneurons. Optotagging experiments showed that these neurons correspond to either PV+ or narrow-waveform Sst+ neurons. A computational model could explain the observed differences in phase locking based on the neurons’ capacitative low-pass filtering properties. In summary, the propagation of synchronized activity and its effect on distinct cell types strongly depend on its frequency.

## Introduction

Rhythmic flicker stimulation has gained increased interest in the context of therapeutic methods for neurodegenerative diseases and as a method for tracking neuronal processes. For instance, recent studies have shown that 40 Hz visual flicker stimulation can reduce beta-amyloid plaques in Alzheimer’s disease mouse models (Singer, 2018; Adaikkan and Tsai, 2020). It has been suggested that these effects of high-frequency rhythmic stimulation depend on brain-wide entrainment including the hippocampus and media prefrontal cortex (mPFC) (Adaikkan et al., 2019). Moreover, high-frequency rhythmic stimulation is increasingly used as a technique for human MEG and EEG recordings to track neural processing across various stages through “frequency tagging” (Zhigalov et al., 2019; Vialatte et al., 2010; Drijvers et al., 2021; Seijdel et al., 2022). Thus, it is important to determine the neural mechanisms through which flicker-induced synchronization propagates across cortical levels, and how flicker stimuli influence distinct cell types in the local circuit.

Synchronized activity can result from either rhythmic external inputs or endogeneous network interactions. Several theoretical and empirical studies suggest that synchronized activity may facilitate the processing of information, by enhancing the impact of spikes on post-synaptic targets (Bernander et al., 1994; Salinas and Sejnowski, 2001; Fries et al., 2001). Synchronization of neural responses may be especially critical for information flow in the case of sparse feedforward connections, e.g. in case of thalamocortical communication (Bruno and Sakmann, 2006). Furthermore, functional studies on inter-areal communication suggest that feedforward influences are particularly strong for high-frequency rhythmic activity (van Kerkoerle et al., 2014; Bastos et al., 2015).

Yet, several observations suggest that high-frequency synchronization might not be conductive to signal propagation: First, it is evident that there is perceptual filtering of flicker stimuli above a certain frequency (flicker fusion threshold). One possibility is that flicker-induced synchronization does effectively propagate across various stages of the cortical hierarchy, but is perceptually filtered out due to other cortical mechanisms (e.g. in higher processing stages). However, studies of mass population activity in humans (magnetoencephalography, MEG) suggest that high-frequency flicker stimuli may not effectively propagate beyond the primary visual cortex (Duecker et al., 2021). Second, low-frequency synchronization is typically seen at a much larger spatial scale than local high-frequency synchronization (Steriade, 2001; Csicsvari et al., 2003; Buzsaki and Draguhn, 2004). Third, the passive integration properties of single neurons related to the cable equation cause dendritic low-pass filtering, especially in pyramidal neurons (Fortune and Rose, 1997; Koch, 2004; Pike et al., 2000a; Vaidya and Johnston, 2013). Although the combination of active and passive integration properties can also create resonance behavior, i.e. the selective amplification of information in a specific frequency range (Hutcheon and Yarom, 2000; Izhikevich et al., 2003; Blankenburg et al., 2015). Thus, it remains overall unclear how synchronization driven by rhythmic stimulation propagates throughout the cortex and how different components of the microcircuit are affected, depending on frequency.

To investigate this, we used Neuropixels to record from multiple processing stages in the mouse brain simultaneously (LGN, different layers of V1, CA1 Hippocampus) while presenting (LED and monitor) flicker stimuli at different frequencies. Using optotagging we distinguished the activity of excitatory neurons and specific GABAergic subtypes, namely PV+ and Sst+ interneurons. To explain our experimental observations, we performed detailed multicompartmental modeling of mouse V1 cells to investigate the filtering properties of the different cell types.

## Results

We recorded isolated single units from areas LGN, V1, and CA1 using Neuropixels probes, while mice were placed on a running disk (Figure 1A, see Methods). Visual flicker stimuli (frequencies between 10 - 80 Hz) were presented using either a monitor or an array of LEDs (Figure 1A-B). In the case of monitor flicker, we flashed full-field black-and-white stimuli. In the case of the LED arrays, we used similar luminance settings as in previous studies that examined the effect of visual flicker on neurodegeneration (Singer et al., 2018). We quantified the locking of individual spikes to the flicker stimuli by first extracting the phase of each spike relative to the flicker cycle and then computing the pairwise phase consistency (PPC) (Figure 1D). The PPC is a measure of phase locking that is unbiased by spike count or firing rates (Vinck et al., 2012).

**Figure 1:**
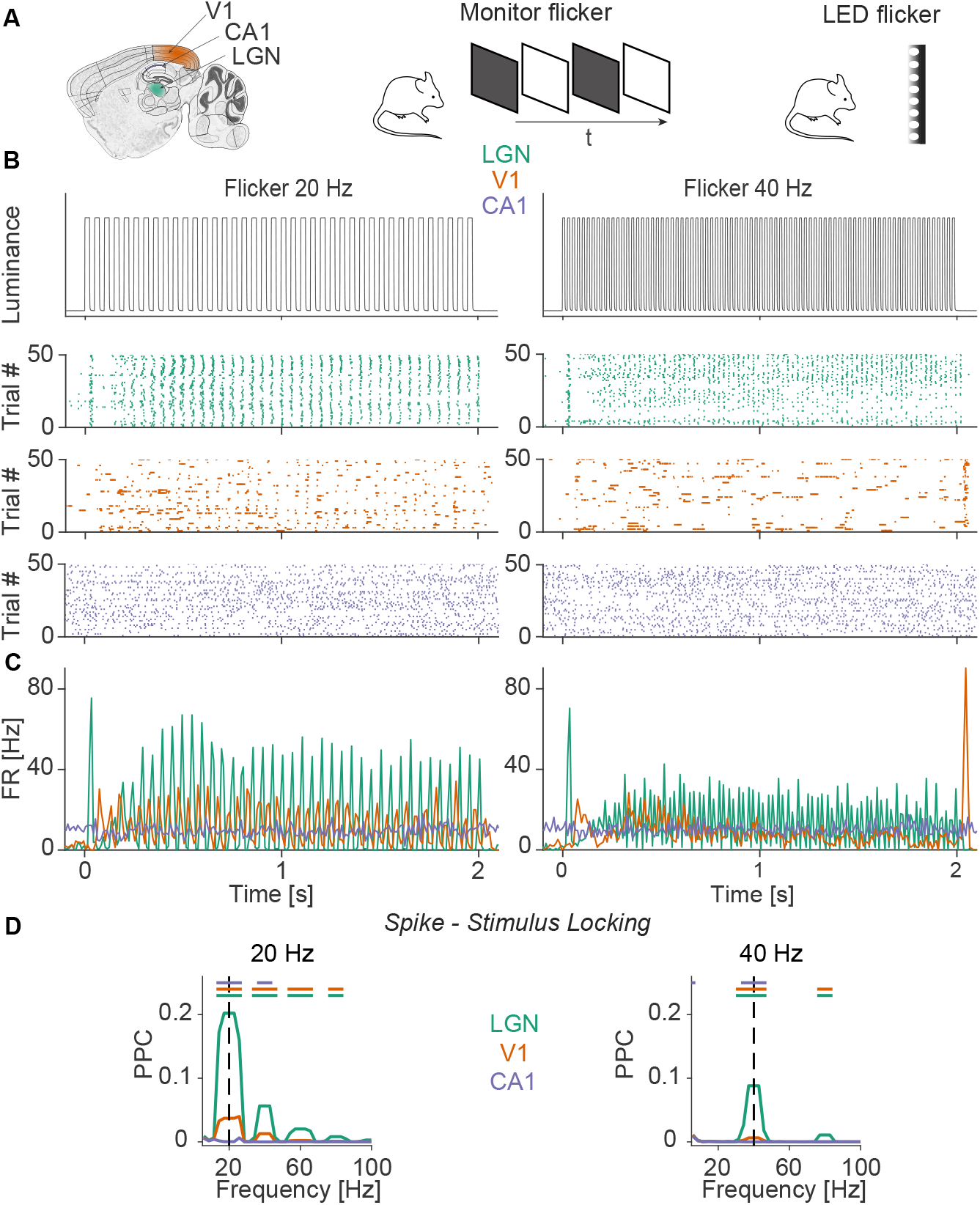
Illustration of the experiment. **(A)** We made simultaneous Neuropixels recordings from LGN, V1, and CA1 while presenting fast flickering stimuli of different frequencies. Flicker frequencies were presented in randomized order. The rhythmic flicker was presented using either a full-screen monitor or LEDs. **(B)** Measures luminance change (using photodiode) of the LED for 20 Hz (top, left) and 40 Hz (top, right) flicker frequency (y-axis has arbitrary units). Raster plot of example neurons in LGN (green), V1 (orange), and CA1 (blue). **(C)** PSTHs of neurons shown in (B). **(D)** Phase-locking of neurons in LGN (N=2386), V1 (N=2091), and CA1 (N=636) to the flicker stimulus. Colored lines on top indicate significantly different frequency bins, non-parametric permutation test, and FDR correction for multiple comparisons with a threshold of P<0.05. Green line: LGN-V1, orange line: LGN-CA1, blue line: V1-CA1. Unless otherwise stated, statistical tests in the study were non-parametric, two-sided, and based on 1000 randomizations.

Significant phase-locking to the flicker stimuli was observed in areas LGN and V1, but not in CA1 (Figure 1B-C for example neurons, Figure 2A for population analysis). For all frequencies, phase locking was stronger in the LGN than in V1 (Figure 2A, see Figure 1D for PPC spectra). Both in V1 and LGN, phase locking decreased with frequency (Figure 2A). Around 40 Hz, LGN units still exhibited significant phase locking to the flicker stimuli, whereas phase locking was an order of magnitude weaker in area V1 (Figure 2B). At 60 Hz and beyond, we did not observe significant phase locking in any of the areas (Figure 2A). Restricting our analysis to visually responsive neurons did not change our results (Figure S1). Similar phaselocking patterns were observed for LED and monitor flicker stimuli (Figure 2B, Figure S2A,B for summary plots of monitor stimuli, S3A for PPC spectra during monitor stimulation).

**Figure 2:**
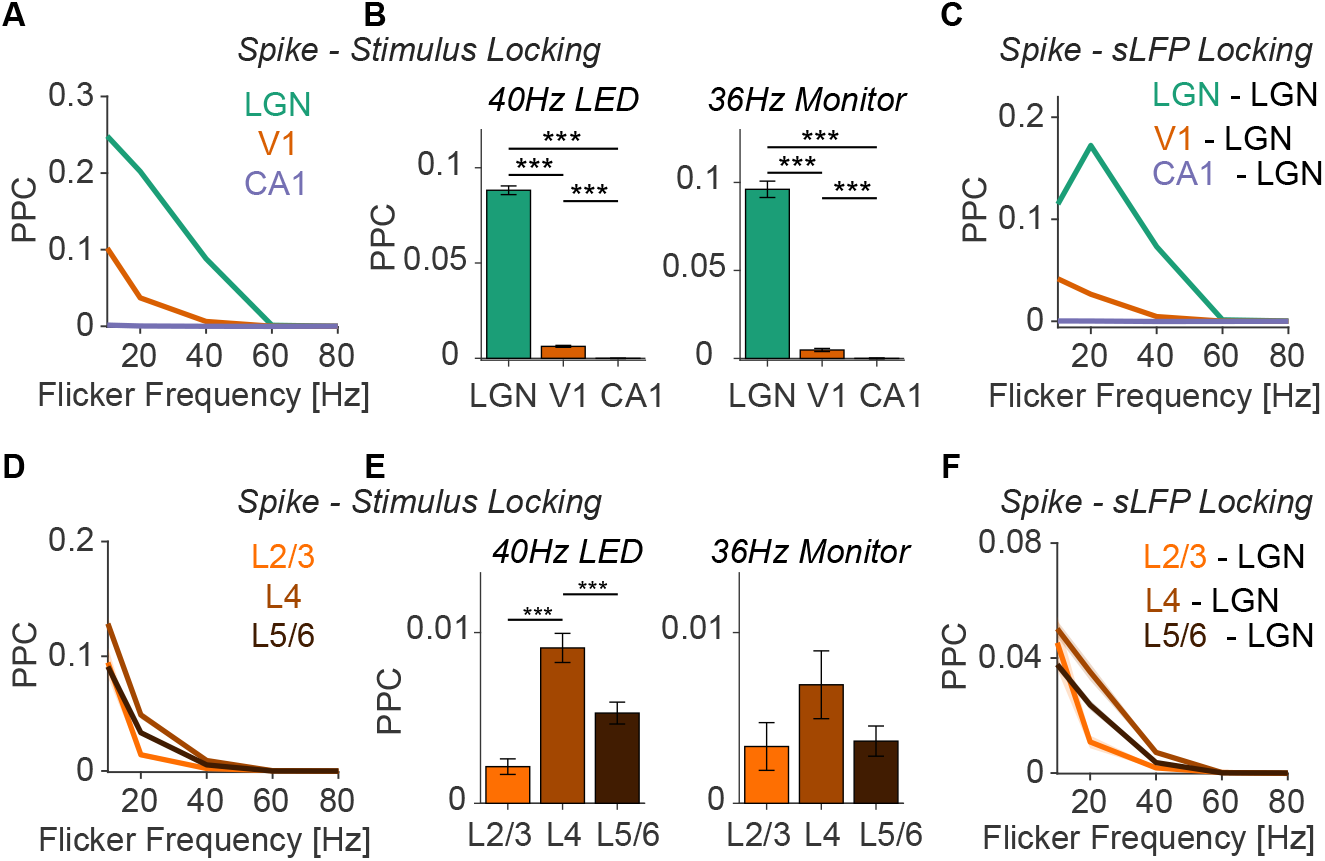
Propagation of flicker-induced synchronization across processing stages. **(A)** Spike-stimulus phase-locking of neurons in LGN (N=2386), V1 (N=2091), and CA1 (N=636) during LED flicker stimulation. Phase locking was measures with the unbiased pairwise phase consistency (PPC, see Methods). **(B)** Phaselocking to stimulus during 40Hz LED (left, N_LGN_ = 2386, N_V1_ = 2091, N_CA1_ = 636) and 36Hz monitor (right, N_LGN_ = 1153, N_V1_ = 815, N_CA1_ = 125) flicker presentation. ***P<0.001; **P<0.01; *P<0.05, non-parametric permutation tests, based on 1000 randomizations. **(C)** Spike-sLFP phase-locking for different flicker frequencies (presented using LEDs) and combinations of spikes and sLFPs: Spikes in LGN to sLFPs in LGN, spikes in V1 to sLFP in LGN, and spikes in CA1 to sLFP in LGN. The sLFP is the surrogate LFP constructed by summing all LGN spikes together and low-pass filtering (see Methods). **(D)** Spike-stimulus phase-locking of neurons in different layers of V1 during LED flicker stimulation N_sup._ = 80, N_gra._ = 604, N_inf._=1407). **(E)** PPC between neurons in different V1 layers and the stimulus during 40HZ LED (left, N_sup._=80, N_gra._=604, N_inf._=1407) and 36Hz Monitor (left, N_sup._=32, N_gra._=279, N_inf._=504) stimulus presentation. **(F)** Spike-sLFP phase-locking during LED flicker presentation between spikes in different V1 layers and the LGN sLFP.

To analyze the phase locking of V1 neurons across cortical layers, we identified different layers using CSD analysis (see Methods). Neurons in the input Layer 4 showed the strongest phase locking to the flicker stimuli, both for monitor and LED flicker (Figure 2D, Figure S2C). Locking was substantially weaker in L2/3, esp. for the 40 Hz LED flicker stimuli (Figure 2E).

It is possible that network synchronization may not have occurred exactly at the frequencies of the flicker stimuli. We therefore also quantified the phase locking of single units to the population activity in the LGN. Because of the geometric arrangement of excitatory cells (closed field), the LGN does not necessarily produce an informative and strictly local LFP. We therefore constructed a “surrogate LFP” (sLFP) for the LGN by summing all LGN spikes (Okun et al. 2015; Schneider et al. 2021, see Methods). Phase locking was then computed as the PPC between spikes and the sLFP. We found that phase locking to the LGN-sLFP showed similar differences between areas and frequencies as compared to phase locking to the flicker stimuli (Figure 2C,F, see Figure S2B,D for phase locking during monitor stimulation). Furthermore, the phaselocking spectra showed narrow-band peaks at the frequencies of the flicker, showing that synchronization was indeed restricted to the flicker frequency (Figure S3B,C).

Together, these analyses indicate that at gamma frequencies, phase locking shows a substantial decrease from LGN to the input L4 of V1 and then further decreases towards L2/3, with no phase locking observed at higher levels of the cortical hierarchy (CA1).

### Fast stimuli primarily recruit fast-spiking interneurons

To analyze the responses of distinct V1 cell types to flicker stimuli, we distinguished cell types based on action potential waveforms (Figure 3) and in a subsequent figure validated these findings using optotagging (Figure 4). Consistent with previous work, V1 neurons were clearly divided into two categories having broad (BW) and narrow (NW) waveforms (Figure 3A). These categories correspond to putative excitatory neurons and fast-spiking interneurons, respectively (Senzai et al. 2019; see also Figure 4). At low frequencies, BW and NW neurons showed approximately similar phase-locking values (Figure 3B-C). However, for higher frequencies, NW neurons were significantly more phase-locked than BW neurons (Figure 3B-D). At gamma frequencies, the strongest phase locking was observed for L4 NW neurons, and phase locking was very weak in L2/3 excitatory neurons (Figure 3D). Similar findings were made for LED and monitor flicker (Figure 3B-D).

**Figure 3:**
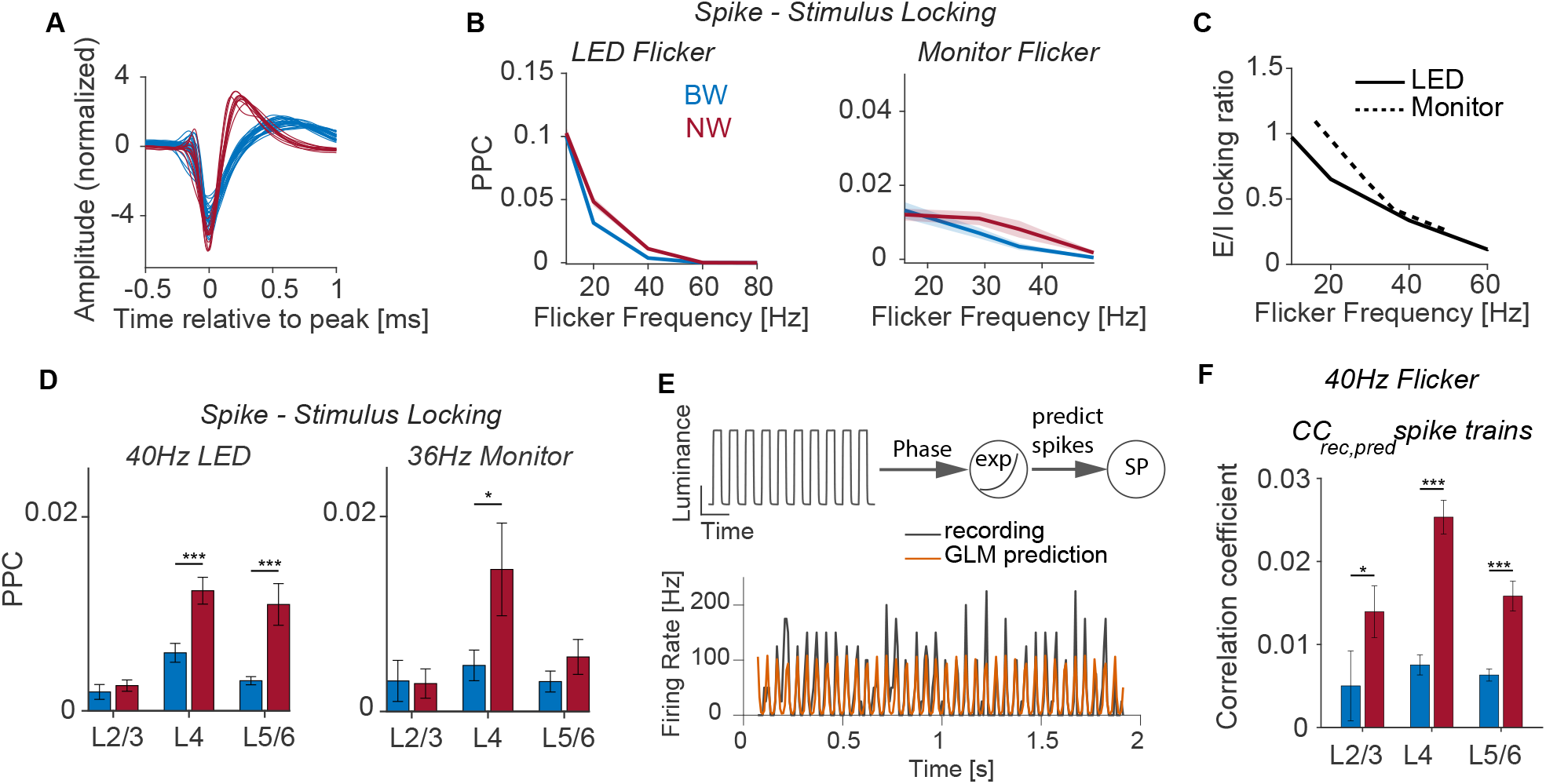
Fast frequencies predominantly drive narrow-waveform interneurons. **(A)** Normalized spike waveforms of neurons in V1. **(B)** Phase-locking between different cell classes in V1 and the LED (left, N_BW_=1316, N_NW_=664) and monitor (right, N_BW_=474, N_NW_=274) during flicker stimulation at different frequencies. **(C)** The ratio between phase-locking of excitatory and inhibitory cells to the stimulus using LEDs (black) and monitor (green). **(D)** Phase-locking of different cell classes in the different layers of V1 to 40Hz LED (left, BW: N_sup._ = 38, N_gra._ = 300, N_inf._ = 978, NW: N_sup._ = 37, N_gra._ = 269, N_inf._ = 358) and 36Hz monitor (right, BW: N_sup._ = 11, N_gra._ = 182, N_inf._ = 375, NW: N_sup._ = 21, N_gra._ = 104, N_inf._ = 109) stimulus. **(E)** Illustration of statistical model employed to predict spike times of V1 neurons from the instantaneous stimulus phase (top). PSTH of recorded example neuron in V1 during 20Hz LED flicker stimulation (black) and corresponding prediction from the model (orange). **(F)** The correlation coefficient between recorded and predicted spike trains of neurons in different layers of V1. (*P<0.05; **P<0.01; ***P<0.001; Wilcoxon Mann-Whitney test). **(D)** N=887 N=535

**Figure 4:**
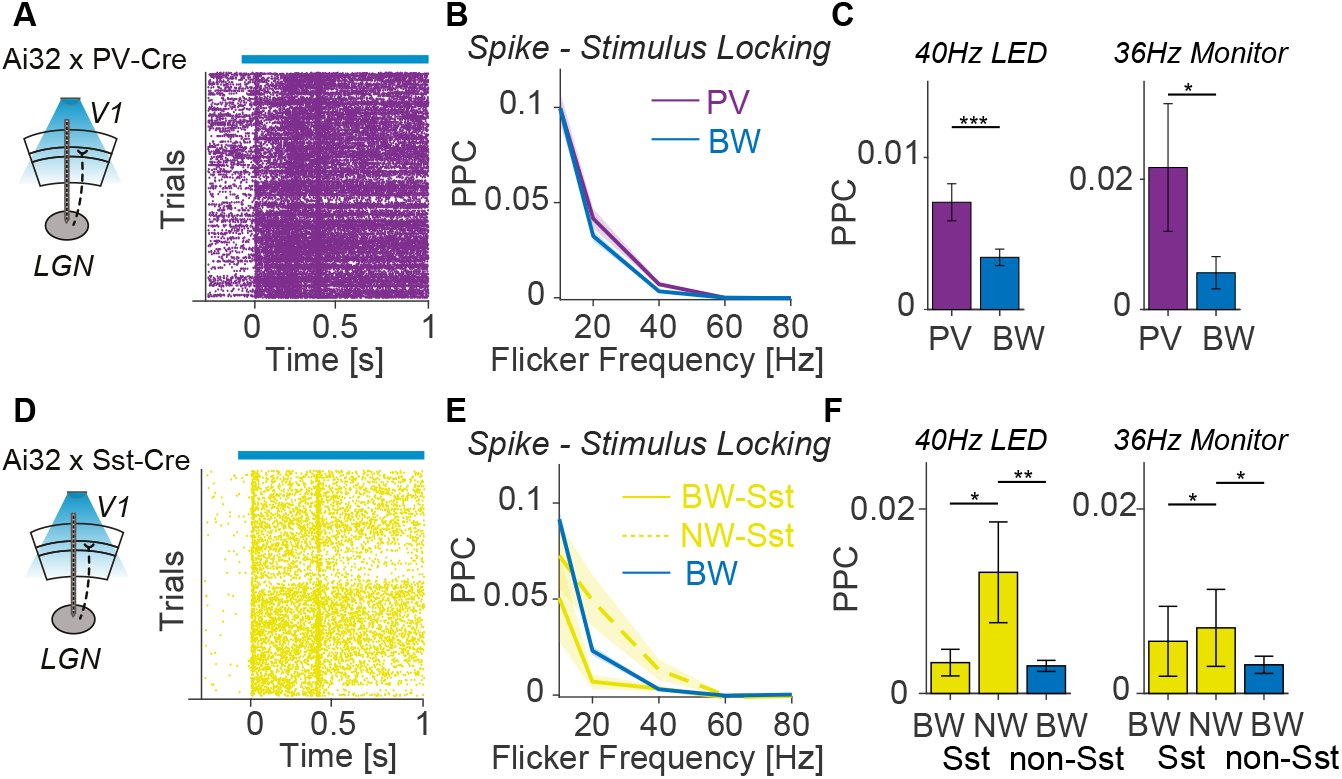
Spike-stimulus phase-locking of GABAergic sub-types. **(A)** Illustration of the experimental setup in PValb-IRES-Cre mice. PV+ cells were identified by their modulation of firing during the stimulation of V1 using a blue laser (see Methods). **(B)** PPC between spikes of PV+ (N=152) as well as BW (N=525) neurons and the LED flicker stimulus. **(C)** Phase-locking of PV+ and BW neurons during 40Hz LED (left, N_PV+_=152, N_BW_=525) and 36Hz monitor (right, N_PV+_=80, N_BW_=104) stimulation. **(D)** Illustration of the experimental setup in SOM-IRES-Cre mice. Sst+ cells were identified by their modulation in firing during the stimulation of V1 using a blue laser (see Methods). **(E)** PPC between spikes of NW Sst+, BW Sst+, as well as BW neurons and the LED flicker stimulus. **(F)** Phase-locking of Sst+ and BW neurons during 40Hz LED (left, N_BW,Sst_=13, N_NW,Sst_=32, N_BW_=541) and 36Hz monitor (right, N_BW,Sst_=8, N_NW,Sst_=26, N_BW_=464) stimulation.

We used a predictive regression model to analyze to what extent the spike times of NW and BW neurons could be predicted by the phase of the flicker stimuli (Figure 3E; see Methods). Consistent with the phase-locking analysis, the spike timing could be substantially better predicted for NW than for BW neurons (Figure 3F). Optotagging experiments were performed in Ai32 x PV-Cre, and Ai32 x Sst-Cre animals to further distinguish between different GABAergic subtypes (Figure 4A,D). Consistent with the analysis of NW neurons, we found that PV+ neurons (which typically had narrow waveforms) were more strongly locked to gamma-frequency flicker stimuli than BW neurons (Figure 4B-C). For Sst+ neurons, we analyzed NW and BW Sst+ neurons separately. At gamma frequencies, NW Sst+ neurons exhibited substantial phase locking to visual flicker stimuli, whereas broad-waveform Sst+ neurons were relatively weakly locked (Figure 4D-E).

### Filtering properties of V1 principle cells

We wondered if the observed differences between cell types could be explained by their respective biophysical properties. To this end, we built detailed biophysical multi-compartmental models from the Allen Institute to test the filtering properties of different V1 cell types. The multi-comparmental models contain a set of 10 active membrane conductances placed in the soma and detailed reconstruted morphologies. Model parameters were optimized to reproduce the firing behavior during somatic whole-cell patch clamp recordings in slices (Gouwens et al. 2018; Arkhipov et al. 2018; Billeh et al. 2020; see Methods).

We first tested the capacitive filtering effects of dendrites of different cell types during the passive propagation of signals from a dendritic arbor to the soma. To this end, we injected sinusoidal currents of different frequencies in dendritic compartments 150 µm from the soma in both pyramidal and PV+ neuron models (Figure 5A). The neuron’s response was examined via the voltage fluctuations in the soma, and the transfer impedance was computed for each stimulation frequency (Figure 5B, Vaidya and Johnston 2013). We found that for low frequencies, pyramidal cells had a higher transfer impedance than PV+ neurons. As the stimulation frequency increased, the transfer impedance of PV+ neurons exceeded the one of pyramidal cells (Figure 5C-D). Reducing the membrane capacitance of pyramidal cell dendrites resulted in an increase in transfer impedance (Figure S5).

**Figure 5:**
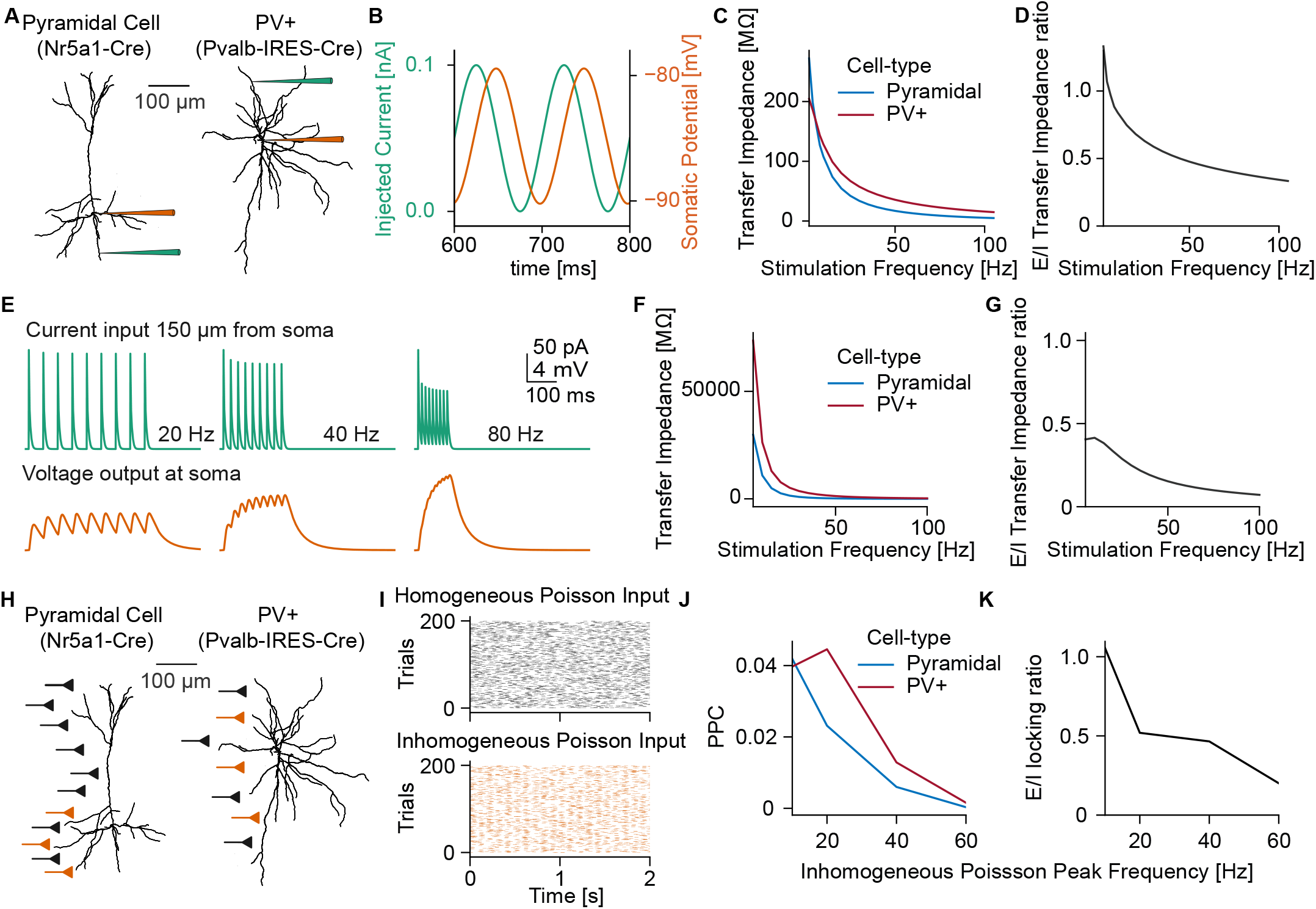
Filtering properties of V1 multi-compartmental neuron models. **(A)** Illustration of Current-Clamp experiment on N5a1-Cre pyramidal neuron and PVAL-IRES-CRE neuron. **(B)** Somatic voltage fluctuations (orange) during sinusoidal current injection into the dendrite at 150 µm (green). **(C)** Transfer impedance of PV+ and Pyramidal cell model during sinusoidal stimulation between 1 and 100Hz. **(D)** Ratio between the transfer impedance of pyramidal (excitatory) and PV+ (inhibitory) neurons. **(E)** Synaptic input current 150 µm from the soma (green, top) and somatic voltage response (orange, bottom). **(F)** Transfer impedance of PV+ and Pyramidal cell model during synaptic burst stimulation between 1 and 100 Hz. **(G)** Ratio between the transfer impedance of pyramidal (excitatory) and PV+ (inhibitory) neurons during synaptic burst stimulation. **(H)** Illustration of synaptic stimulation with homogeneous and inhomogeneous Poisson input. **(I)** Raster plot of example neurons from homogeneous (black) and inhomogeneous (orange) Poisson spiking input population. **(J)** Phase-locking of Pyramidal (blue) and PV+ (red) to sLFP of inhomogeneous Poisson spiking input population. **(K)** Ratio between the phase locking of pyramidal (excitatory) and PV+ (inhibitory) cell models.

In addition, we examined how synaptic bursts with different inter-spike-intervals arriving at a dendrite translate to voltage fluctuations in the soma (Figure 5E). Specifically, we placed one synapse at a dendrite 150 µm from the soma and varied the frequency of the synaptic input between 1 and 100 Hz. We found that the transfer impedance during synaptic burst stimulation was in general higher for PV+ than for pyramidal cells, especially for higher frequencies (Figure 5F-G).

To quantify the phase locking of the neurons’ output spikes to the rhythmic input, we performed simulations that incorporated the active channel properties contributing to spiking outputs. In these simulations, we placed two sets of synaptic inputs at the dendrites of the different cell types. A first subset of synapses, placed along the whole dendritic tree, were activated using homogeneous Poisson spike trains, which mimicked the uncorrelated background activity of V1 cells in the flickering experiment. In addition, we placed a second set of synapses at the basal dendrites of pyramidal cells and along the entire dendritic tree of PV+ cells. The latter synapses were activated by an inhomogeneous Poisson process (modulated at frequencies between 10 and 80 Hz) to simulate input from the LGN to V1 during visual flicker stimulation (Figure 5H,I). The modulation strength of the inhomogeneous Poisson input spike-trains was adjusted to match the phase locking of the recorded neurons in LGN during 10 Hz flicker stimulation (Figure S4A). The number of synapses and synaptic weights was fitted to reproduce the firing rates and phase locking of the two neuron classes during 10 Hz LED stimuli (Figure S4B; see Methods). After fitting the connectivity parameters based on the 10 Hz stimulation, we generated inhomogeneous spike trains modulated at higher frequencies (20 Hz, 40 Hz, 60 Hz and 80 Hz). The modulation strength of the inhomogeneous input spike trains was adjusted to reproduce the observed spike sLFP phase locking in the LGN recordings (Figure S4C. Increasing the frequency of the input rhythm resulted in a substantial decrease in the phase locking of neurons. This decrease in phase locking with increasing stimulation frequency was much steeper in pyramidal than PV+ neurons, similar to the experimental findings reported above. Increasing the dendritic membrane capacitance in pyramidal cells resulted in stronger phase locking at high frequencies, enabling pyramidal cells to follow a 40 Hz stimulation similar to PV+ cells (Figure S5C-D).

## Discussion

We investigated the propagation of rhythmic visual flicker stimulation throughout different stages of the visual hierarchy. Strong phase-locked responses in LGN neurons were induced by presenting flickering stimuli between 10 and 80 Hz using either LEDs or a monitor. LGN neurons showed strong phase locking at flicker frequencies up to 40 Hz, whereas phase locking was substantially weaker in V1 units and absent in hippocampal CA1 units. Separating neurons in the different layers of V1 revealed an attenuation of phase locking at each processing stage of V1, esp. at gamma frequencies. Phase locking was strongest in the input layer and substantially weaker phase locking in the superficial layers. Stimuli flickering in the gamma-frequency range predominantly caused phase-locking in neurons with a narrow action-potential waveform, which is characteristic of fast-spiking interneurons. Optogenetic-tagging experiments showed that PV+ and NW Sst+ exhibit substantially stronger phase locking to high-frequency stimuli as compared to excitatory BW neurons or BW Sst+ neurons. Finally, a computational model could explain the observed differences in phase locking based on the neurons’ capacitative low-pass filtering properties.

### Propagation of flicker-induced synchrony

Our findings have several implications for the use of frequency tagging as a method to track neural processing. While most studies on frequency tagging used low-frequency stimuli, recent studies suggest that using higher frequencies than the flicker fusion threshold may have two advantages: First, the use of high frequencies avoids interference of the consciously perceived flicker with the task (Seijdel et al., 2022). Second, using high frequencies may avoid interference of the rhythmic stimuli with endogenous oscillations (Seijdel et al., 2022). However, in the present study, we showed a clear disadvantage of using higher frequencies for frequency tagging: We observed little propagation of high-frequency synchronization beyond the granular input layer of V1 and a reduction in phase locking of about an order of magnitude per processing stage. Thus, using high frequencies might be primarily useful for tagging neural activity and studying excitability in early processing stages. This conclusion is in line with recent human EEG and MEG studies showing that strong responses at the tagged frequency are mostly restricted to early sensory areas (Williams et al., 2004; Zhigalov et al., 2019; Drijvers et al., 2021; Duecker et al., 2021; Marshall et al., 2022).

A key difference between our study and human studies of frequency tagging is that we examined single-neuron spiking activity. Analysis of field potential (e.g. MEG, EEG) signals cannot determine whether spiking activity is phase-locked to flicker activity. The reason is that field potential signals reflect synaptic inputs that derive from two potential sources: (1) afferent inputs from other areas and (2) local spiking activity (Schneider et al., 2021; Buzsáki and Schomburg, 2015; Pesaran et al., 2018). Thus, MEG/EEG signals in V1 can be generated by spatiotemporally coherent LGN afferents driving EPSPs in the V1 input layer. The extent to which MEG/EEG signals reflect local spiking activity depends on the extent to which local neurons are phase-locked to the afferent inputs. The coherence of MEG/EEG signals with the flicker stimulus may thus reflect thalamocortical inputs. Our results further suggest that the relative contribution of cortical spiking and thalamocortical afferents to MEG/EEG signals depends on the flicker frequency: At lower frequencies, V1 phase-locking is relatively strong, and MEG/EEG signals should therefore reflect approximately equal contributions from afferent inputs and local spiking activity (Schneider et al., 2021). By contrast, at higher frequencies, V1 phase-locking becomes very weak as compared to LGN phase-locking, which predicts that the coherence of EEG/MEG signals with the flicker stimulus is largely driven by thalamocortical afferents.

Recent studies have shown a significant decrease in Alzheimer-associated beta-amyloid plugs across many cortical regions (V1, PFC, CA1, S1) following chronic 40 Hz LED flicker stimulation (Iaccarino et al., 2016; Singer, 2018; Adaikkan et al., 2019). These studies also showed significant LFP-LFP coherence between the visual cortex and higher cortical regions including PFC and CA1, which was suggested as a potential mechanism underlying the effects of flicker on neurodegeneration (Adaikkan et al., 2019). Here, we did not observe phase locking of single CA1 neurons to the flicker stimulus, which is unlikely due to a lack of sensitivity as we recorded from a large sample of CA1 neurons. Furthermore, we observed a strong attenuation of high-frequency synchronization already at early processing stages. This raises the question, as to whether the previously observed V1-CA1 LFP-LFP coherence in fact reflects the phase locking of CA1 spikes. Considering that the visual cortex is located in close proximity to CA1, it is generally difficult to rule out volume conduction (Sirota et al., 2008; Vinck and Bosman, 2016). Another key difference between our study and the previous neurodegeneration study is that we did not use chronic flicker stimulation across several weeks. It is possible that long-term chronic flicker stimulation induces synaptic plasticity which could lead to an enhancement of entrainment in higher-order brain regions across weeks. Nevertheless, it remains a possibility that the therapeutic effects of flicker stimulation on neurodegeneration in CA1 are not mediated by the phase locking of CA1 neurons.

### Implications for endogenous synchronization

Our observations of weak propagation of high-frequency synchronization are in line with our recent findings in macaque and mice: Spyropoulos et al. (2022) find that endogenous gamma-frequency oscillations in awake macaque V1 mainly recruit V4 fast-spiking inhibitory interneurons that reside in the granular input layer of V4, with no phase locking in superficial layers of V4. Similar observations were made for the propagation of LGN gamma synchronization to area V1 in mice Spyropoulos et al. (2022). As in the present study, optotagging experiments in mice further suggest that endogenous LGN gamma predominantly drives fast-spiking PV+ and Sst+ interneurons in the input layer of V1 Spyropoulos et al. (2022). The observation that CA3 gamma synchronization primarily recruits CA1 interneurons (Schomburg et al., 2014) suggests a motif that may be prevalent across many cortical regions. The computational models presented here suggest that the preferential recruitment of fast-spiking interneurons in the granular layer by high-frequency afferents can be explained by differences in capacitive low-pass filtering properties. This conclusion is in line with previous studies that have examined single-neuron filtering properties in fast-spiking interneurons and excitatory neurons (Pike et al., 2000b; Izhikevich et al., 2003; Hasenstaub et al., 2005).

The weak propagation of high-frequency synchronization and its effect on different cell types contradicts the hypothesis that gamma synchronization promotes feedforward information transmission (Bastos et al., 2015; Fries, 2015; Salinas and Sejnowski, 2001). We note that the evidence for this hypothesis is exclusively based on the analysis of LFP-LFP Granger-causality (van Kerkoerle et al., 2014; Bastos et al., 2015), which can therefore not determine the effect of synaptic afferents on spiking activity in the downstream receiver (Schneider et al., 2021). However, our findings match with several observations: First, the observation that high-frequency oscillations are mainly locally coherent and recruit neurons mostly restricted to the area of origin, in contrast to more globally coherent slower rhythms (Steriade, 2001; Csicsvari et al., 2003; Buzsaki and Draguhn, 2004; Buzsáki and da Silva, 2012). Second, the observation that strong (endogenous) gamma synchronization in LGN and primary visual cortex is mostly observed during the presentation of highly predictable, low-dimensional stimuli which have on average a low salience (Schneider et al., 2021; Uran et al., 2022; Vinck et al., 2022).

### Conclusions

We show here how synchronized activity, induced by flicker stimuli, propagates across brain areas and affects distinct cell types depending on the frequency. Specifically, our findings suggest that low-frequency synchronization propagates effectively across multiple cortical stages and recruits excitatory and inhibitory neurons to a similar extent, whereas high-frequency synchronization tends to stay local and primarily recruits fastspiking interneurons downstream. This could suggest that when a neural population switches from low to high-frequency synchronization, output signals will be differently integrated by a downstream receiver. These findings have several implications for the understanding of frequency tagging methods and the mechanisms underlying the effects of high-frequency stimulation on neurodegeneration.

## Authorship contributions

Conceptualization: MS and MV. Simulations and data analysis: MS. Mouse recordings and surgeries: AT. Software and methods for mouse data: MS, AT, CU, MV. Writing: MS and MV. Supervision: MV. This project was supported by ERC Starting Grant to MV (SPATEMP) and a BMF Grant to MV (Bundesministerium fuer Bildung und Forschung, Computational Life Sciences, project BINDA, 031L0167).

## Methods

### LEAD CONTACT AND MATERIALS AVAILABILITY

The Lead Contact of this study is Martin Vinck. Further information and requests for resources should be directed to and will be fulfilled by the Lead Contact, Martin Vinck.

### DATA AND CODE AVAILABILITY

The open-source MATLAB toolbox Fieldtrip (Oostenveld et al., 2011) was used for data analysis. Data and custom MATLAB scripts are available upon request from Martin Vinck.

### EXPERIMENTAL MODEL AND SUBJECT DETAILS

Experiments were performed on three to eight months old male mice. All procedures complied with the European Communities Council Directive 2010/63/EC and the German Law for Protection of Animals and were approved by local authorities, following appropriate ethics review. Mice were maintained on a 12/12 h light/dark cycle and recordings were performed during their dark (awake) cycle. To identify the PV-positive (PV+) and SST-positive neurons (SST+) during electrophysiological recordings, we crossed PV-Cre-mice (B6.129P2-Pvalbtm1(cre)Arbr/J, JAX Stock 017320, The Jackson Labaratory) to Ai32(RCL-ChR2(H134R)/EYFP), JAX Stock 024109, The Jackson Labaratory) mice, and Sst-IRESCre mice (Ssttm2.1(cre)Zjh, JAX Stock 013044, The Jackson Labaratory) to Ai32(RCL-ChR2(H134R)/EYFP) mice, to allow Cre-dependent expression of ChR2 in PV+ (PV-ChR2) and SST+ neurons (SST-ChR2), respectively.

#### Flicker stimulation

In the first set of experiments, visual flicker stimuli were generated using Psychophysics Toolbox (Brainard and Vision, 1997). The experiment was run on a Windows 10 and stimuli were presented on a monitor with a 144 Hz refresh rate. Square wave flicker stimulation was presented on the full screen at 4 different frequencies (16 Hz, 29 Hz, 36 Hz, and 49 Hz). In the second set of experiments, visual flicker stimuli were presented using light-emitting diodes (LEDs) with 5 different pulse frequencies (10 Hz, 20 Hz, 40 Hz, 60 Hz, and 80 Hz). Visual flicker stimulation was generated as described (Singer, 2018) with matching LEDs and other components. The array of LEDs was placed in front of the head-fixed mice at a distance of 17 cm emitting square wave pulses. LEDs had a correlated color temperature (CCT) of 4,000 K and an intensity of 200 lux at the head-post position measured using a Flame UV-VIS Miniature Spectrometer. The trial length for each frequency was 2s with randomized inter-stimulus intervals of 4-10 s.

### METHOD DETAILS

#### Neuropixels recordings and optogenetics

All procedures complied with the European Communities Council Directive 2010/63/EC and the German Law for Protection of Animals and were approved by local authorities, following appropriate ethics review. To identify the PV-positive (PV+) and SST-positive neurons (SST+) during electrophysiological recordings, we crossed PV-Cre-mice (B6.129P2-Pvalbtm1(cre)Arbr/J, JAX Stock 017320, The Jackson Laboratory) to Ai32(RCL-ChR2(H134R)/EYFP), JAX Stock 024109, The Jackson Laboratory) mice, and Sst-IRES-Cre mice (Ssttm2.1(cre)Zjh, JAX Stock 013044, The Jackson Laboratory) to Ai32(RCL-ChR2(H134R)/EYFP) mice, to allow Cre-dependent expression of ChR2 in PV+ (PV-ChR2) and SST+ neurons (SST-ChR2), respectively. Experiments were performed in the three to eight months old male mice. Thirty minutes prior to the head-post surgery antibiotic (Enrofloxacin, 10 mg/kg, sc, Bayer, Leverkusen, Germany) and analgesic (Metamizole, 200 mg/kg, sc) were administered. For the anesthesia, induction mice were placed in an induction chamber and briefly exposed to isoflurane (3 % in oxygen, CP-Pharma, Burgdorf, Germany). Shortly after the anesthesia induction, the mice were fixated in a stereotaxic frame (David Kopf Instruments, Tujunga, California, USA) and the anesthesia was adjusted to 0.8 – 1.5 % in oxygen. To prevent corneal damage the eyes were covered with eye ointment (Bepanthen, Bayer, Leverkusen, Germany) during the procedure. A custom-made stainless steel head fixation bar was secured with dental cement (Super-Bond C & B, Sun Medical, Shiga, Japan) exactly above the bregma suture, while the area of the recording craniotomy (V1, AP: 1.2 mm anterior to the anterior border of the transverse sinus, ML: 2.1 to 2.5 mm) was covered with cyanoacrylate glue (Insta-Cure, Bob Smith Industries Inc, Atascadero, CA USA). Four to six days after the surgery, the animals were habituated for at least five days in the experimental conditions. The day before or the same day of the first recording session a 0.8 mm^2^ craniotomy was performed above V1 (AP: 1.2 mm anterior to the anterior border of the transverse sinus, ML: 2.1 to 2.5 mm) under isoflurane anesthesia. The craniotomy was covered with silicon (Kwik-Cast, World Precision Instruments, Sarasota, USA), and the mouse was allowed to recover for at least 2 hours. Recording sessions were carried out daily for a maximum of 5 days, depending on the quality of the electrophysiological signal. Awake mice were head-fixed and placed on the radial wheel apparatus. We recorded simultaneously from 384 recording sites on a single Neuropixels probe, from LGN, CA1 and V1. The probe was coated with the fluorescent dye DiI (D7757, Thermo Fisher Scientific) and was inserted in the brain tissue through the V1 craniotomy under a 15 degrees angle. We targeted PV+ and SST+ interneurons using PV-ChR2 and SST-ChR2 mice and activated them using optogenetic stimulation. During the optogenetic experiment, an optic fiber (Thorlabs, 200um, 0.39 NA) coupled to a diode laser (LuxX CW, 473 nm, 100 mW, Omicron-Laserage Laserprodukte GmbH, Germany) was used to illuminate V1 craniotomy. The optic fiber was positioned 0.2 mm from the probe position, just above the surface of the brain. Continuous light square pulses were applied for 500 ms interleaved by 3-6 s intervals. The light intensity on the tip of the fiber was 0.02 - 50 mW/mm^2^.

Single units were isolated using the semi-automated spike sorting algorithm Kilosort 2.5 (Steinmetz et al., 2021). To obtain LFPs, electrode signals were first low-pass filtered at 400 Hz and then high-pass filtered at 0.1 Hz, using a third-order Butterworth filter. In order to filter out line noise, an additional band-stop filter between 49.5 and 50.5 Hz and 99 and 101 Hz was applied. Subsequently, signals were downsampled to 1200 Hz by averaging consecutive frames. For memory reasons, only every second electrode was used for the analysis of LFP signals. The pairwise phase consistency (PPC) between spikes and stimuli and spikes and LFPs was calculated using windows of 250 ms around each spike (Vinck et al., 2012), using the ft spiketriggeredspectrum functions in the FieldTrip SPIKE toolbox. Only neurons firing at least 150 spikes were considered for the calculation of spike-LFP and spike-stimuli PPC. Because LGN is a nucleus and the neurons are not aligned, the LFP signal in LGN does not reflect the oscillatory activity of the neurons in LGN. For this reason, we used a surrogate LFP (sLFP) derived from the spiking activity of the neurons in LGN (Okun et al., 2015; Schneider et al., 2021). The sLFP was derived by summing the spikes of all individual isolated units in the LGN. Subsequently, the population spike activity was filtered between 1 and 100 Hz.

#### Identification of CA1 pyramidal cell layer

The hippocampal CA1 pyramidal cell layer was identified based on several physiological criteria such as sharp wave ripples, increased single-unit activity, and large waveform amplitudes (Mizuseki et al., 2011; Fernández-Ruiz et al., 2019). LFP signals from each electrode were band-pass filtered between 130 and 200 Hz followerd by a transformation to a normalized squared signal (NSS). Ripple events were identified as peaks beyond 5 SD above the mean of the normalized squared signal, with a duration between 20 ms and 200 ms. The CA1 pyramidal layer was identified as the recording site with the largest mean power during ripple events. The site with the largest spike waveform amplitude and increased spiking activity in proximity to the recording site with the largest mean ripple power was regarded as the site of CA1 pyramidal cell bodies. All physiological localizations were followed by histological verification. The scripts for ripple event detection (bz FindRipples) can be found at the Buzsaki lab GitHub repository https://github.com/buzsakilab/buzcode.

#### Waveform classification

The mean waveform was calculated over data segments from -41 to 42 samples around the time of the spike, based on the aligned waveforms of the first 10000 spikes of each neuron. The sampling rate was increased by a factor of 3 using spline interpolation. The mean waveforms were normalized by subtracting the median of the first 10 samples and then dividing by the absolute value of the negative peak. Waveforms with a positive absolute peak were discarded. Subsequently, two-dimensional t-Stochastic Neighbor Embedding (t-SNE; perplexity of 80) was applied on the 80 samples after the negative spike peak of the waveforms. Lastly, we applied hierarchical clustering on the two-dimensional t-SNE embedding, which resulted in two separate clusters corresponding to the broad and narrow waveform neurons.

#### Assignment of cortical layers in V1

The assignment of superficial, granular, and deep cortical layers in mouse V1 was based on the current source density (CSD) of the average LFP signal during whole screen flash stimulation. The protocol consisted of a 100 ms long white screen period with a 2 s lasting grey-screen inter-stimulus period. To increase the spatial sampling rate, LFP traces were interpolated with an interpolation factor of 4. Current source density analysis was computed by taking the second discrete spatial derivative across the different electrode recordings sites (Mitzdorf, 1985). The stepsize of the discrete spatial derivative was 200 µm. Single units were assigned to a cortical layer based on the location of the channel with the highest amplitude during a spike.

#### Testing optogenetic response

Optogenetic tagging experiments were performed on Pvalb-IRES-Cre and Sst-IRES-Cre knock-in mice. The optogenetic stimulation consisted of 300 trials of 1s long stimulation periods with a randomized interstimulus interval between 4 and 7 seconds. Cells expressing Cre were identified using the Zeta-test (Montijn et al., 2021). The Zeta test is a recently developed parameter-free statistical test that can be used to determine whether neurons show a time-dependent modulation of their firing rates by an event. The Zeta test was applied to the period around the laser-onset (−10 ms, 10 ms) to test which neurons showed significantly modulated spiking activity (P <0.05). Cells were classified as optogenetically tagged if there they exhibited a significant modulation and if their first crossing of peak half-height, occurred within the 10 ms following the onset of the laser. To avoid misclassification due to laser artifacts, neurons with an onset time of above peak earlier than 1 ms after the onset of the optogenetic stimulation were discarded.

#### Testing visual responsive neurons

The visually responsive neurons were identified using the Zeta-test on the protocol for mapping cortical layers in V1. The Zeta test was applied to the period around the onset of the white screen (0 ms,10 ms) to test which neurons showed significantly modulated spiking activity (P<0.05). Cells were classified as visually responsive if their firing rate was significantly modulated by the onset of the white screen.

#### Statistical modeling

We fitted a regression model to predict neural spiking activity *r*_*rec*_(*t*) from the phase ϕ(*t*) of the flickering stimulus using maximum likelihood estimation. The phase of the flickering stimulus was extracted by calculating the wavelet transform of the signal of a photodiode placed in front of the LEDs. The model is given by:

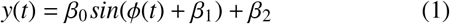

where ϕ(*t*) is the phase of the stimulus, and β_0_, β_1_, and β_2_ are regression parameters. To estimate the firing rate *r*(*t*), the model function *y*(*t*) was passed through an exponential link function.

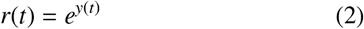

Regression parameters were optimized by minimizing the negative log-likelihood function. The performance of the model was evaluated by computing the Pearson correlation coefficient between the recorded and the predicted spike trains. Spike trains and stimulus traces were downsampled to a sampling rate of 125 Hz. Regression models were fitted using 90 % of the trials of each flicker stimulation frequency and validated based on the 10 % held-out trials.

#### Multi-compartmental models

Simulations were carried out using NEURON (http://www.neuron.yale.edu, Hines 1984; Hines and Carnevale 1997) and the Brain Modeling ToolKit (BMTK) (Dai et al., 2020). The perisomatic models used in this study consist of realistic reconstructions of the dendritic trees and a wide variety of active and passive membrane mechanisms, including 10 types of ion channels placed in the soma (Gouwens et al., 2018). The details on the model and how ion channel parameters were tuned based on electrophysiological recordings can be found on the website of the Allen Institute http://help.brain-map.org/display/celltypes/Documentation. We used two different models of V1 Neurons, Nr5a1 (CellID = 472451419), representing a subclass of L4 and L5 excitatory pyramidal cells in mouse V1 and PV-IRES-Cre (CellID = 471085845), representing a fast-spiking inhibitory cell class in mouse V1 (Gouwens et al., 2018; Nandi et al., 2022).

For Figure 5A-D the simulations ran 1400 ms with time steps of 0.001 ms. Following a preprun of 200 ms, a 1 s long 100 pA sinusoidal current (frequency range: 2-10 Hz, increment 2 Hz and 10 - 105 Hz, increment 5 Hz) was injected in a randomly selected dendrite (in the case of the pyramidal cell model we selected a basal dendrite) at a distance of 150 µm from the soma. Electrical transfer impedance| *Z*(*f*) | was measured as the ratio of the Fourier transform of the membrane voltage in the soma to that of the current input at the dendrite (Vaidya and Johnston, 2013).

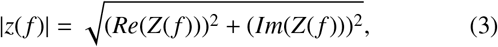

where Re(Z(f)) and Im(Z(f)) are the real and imaginary parts of the ratio of the Fourier transforms at frequency f.

For Figure 5 E-G an excitatory synapse was placed at a randomly selected dendrite (in the case of the pyramidal cell model we selected a basal dendrite) at a distance of 150 µm from the soma. Synaptic stimulation was modeled with an Exp2Syn point process with the following parameters: Synaptic rise time τ_1_ = 1 ms, synaptic decay time τ_2_ = 3 ms, reversal potential E = 0 mV, synaptic delay T = 1 ms, and synaptic weight g_syn_ = 0.004 µS. Synaptic bursts consisted out of 9 spikes spiking with a rate between 5 and 100 Hz (increment 5 Hz).

For Figure 5 H-K we placed two groups of synapses along the dendrites to reproduce the activity recorded during 10 Hz LED flicker stimulation. The first set of synapses was driven by a population of homogeneous Poisson spiking neurons and placed along the whole dendritic tree (basal and apical), simulating the continuous background input through recurrent connections. The second set of synapses, driven via a population of inhomogeneous Poisson spiking neurons (modulated at frequencies 10 Hz, 20 Hz, 40 Hz, and 60 Hz), was placed along the dendrites (for the Nr5a1-Cre model only on the basal dendrites), simulating the rhythmic drive from LGN during visual flicker stimulation. We fitted the number of inhomogeneous and homogeneous spiking neurons terminating on the two cell classes and the synaptic weights in order to reproduce the firing rates and the phase-locking to the inhomogeneous spiking input population of the broad and narrow waveform neurons during 10 Hz LED flicker stimulation. Nr5a1-Cre neurons received inhomogeneous spiking input from 6 neurons, while PV-IRES-Cre received input from 8 inhomogeneous spiking neurons. Nr5a1-Cre neurons received homogeneous spiking input from 60 neurons, while PV-IRES-Cre received input from 43 inhomogeneous spiking neurons. All input neurons had 70 synaptic connections to our model neurons with a synaptic weight of g_syn_ = 0.000012 µS and synaptic delay of T = 4 ms. Synaptic time constants were the same as in Figure 5 E-G. The simulations were run for 10 s with 200 repetitions.

**Figure S1:**
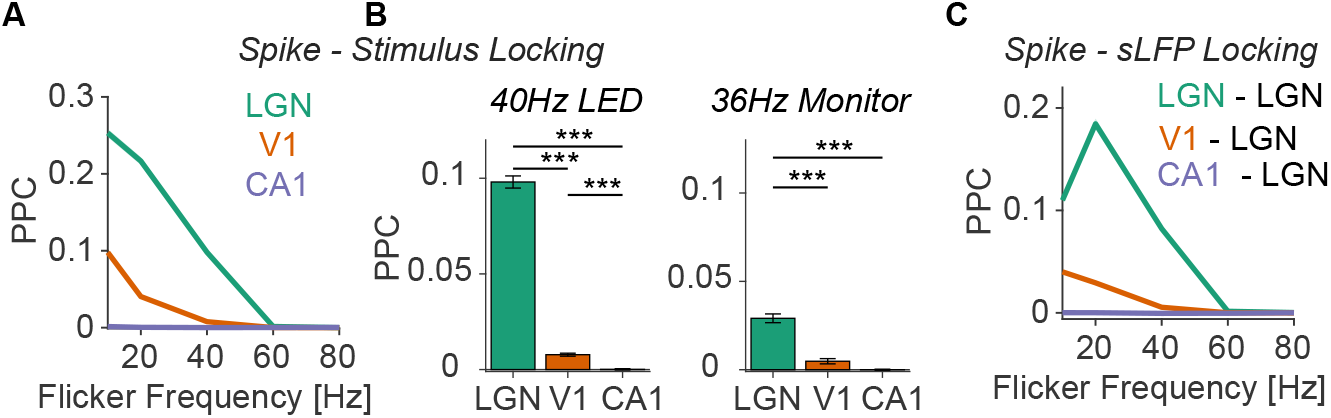
Same as Figure 2 but including only visually responsive neurons. **(A)** Spike-stimulus phase-locking of visually responsive neurons in LGN (N=1332), V1 (N=1028), and CA1 (N=72) during LED flicker stimulation. **(B)** Phase-locking to stimulus during 40Hz LED (left, N_LGN_ = 1332, N_V1_ = 1028, N_CA1_ = 72) and 36Hz monitor (right, N_LGN_ = 461, N_V1_ = 336, N_CA1_ = 16) flicker presentation. ***P<0.001; **P<0.01; *P<0.05, non-parametric permutation tests, based on 1000 randomizations. **(C)** Spike-sLFP phase-locking for different flicker frequencies (presented using a LED) and combinations of spikes and sLFPs: Spikes in LGN to sLFPs in LGN, spikes in V1 to sLFP in LGN, and spikes in CA1 to sLFP in LGN.

**Figure S2:**
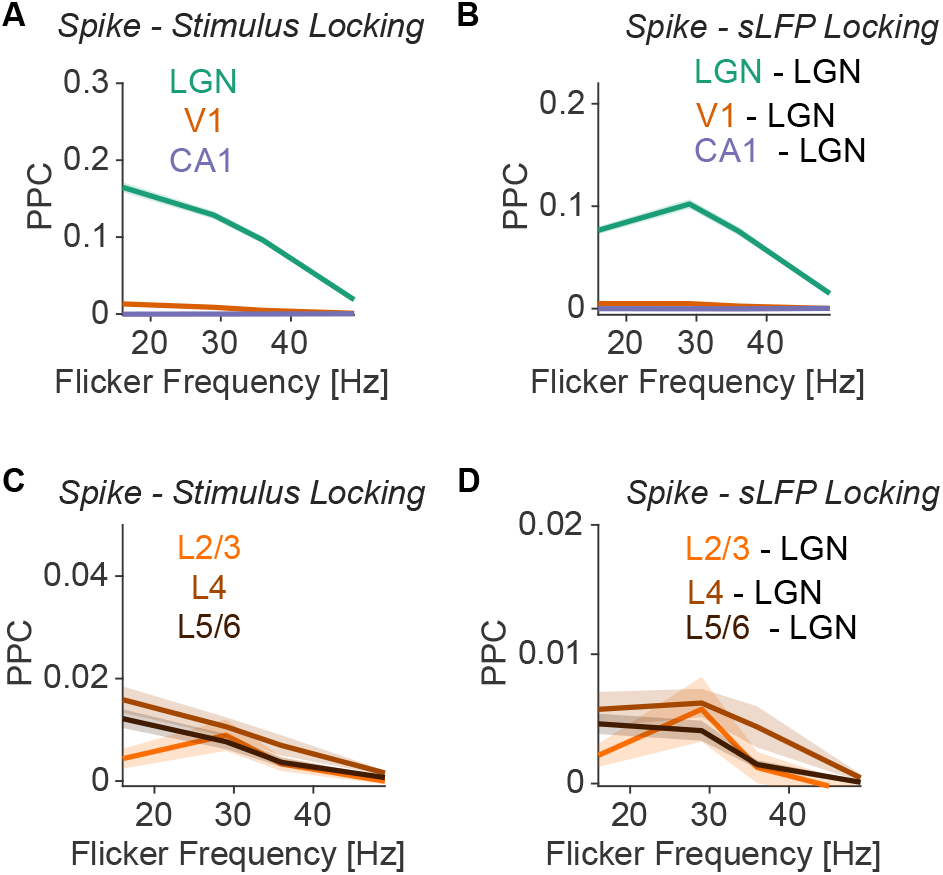
Same as Figure 2 but for monitor stimulation. **(A)** Spike-stimulus phase-locking of neurons in LGN (N=1154), V1 (N=815), and CA1 (N=125) during monitor flicker stimulation. **(B)** Spike-sLFP phase-locking for different flicker frequencies (presented using a monitor) and combinations of spikes and sLFPs: Spikes in LGN to sLFPs in LGN, spikes in V1 to sLFP in LGN, and spikes in CA1 to sLFP in LGN. **(C)** Spike-stimulus phase-locking of neurons in different layers of V1 during monitor flicker stimulation N_sup._ = 32, N_gra._ = 279, N_inf._=504). **(D)** Spike-sLFP phase-locking during monitor flicker presentation between spikes in different V1 layers and the LGN sLFP.

**Figure S3:**
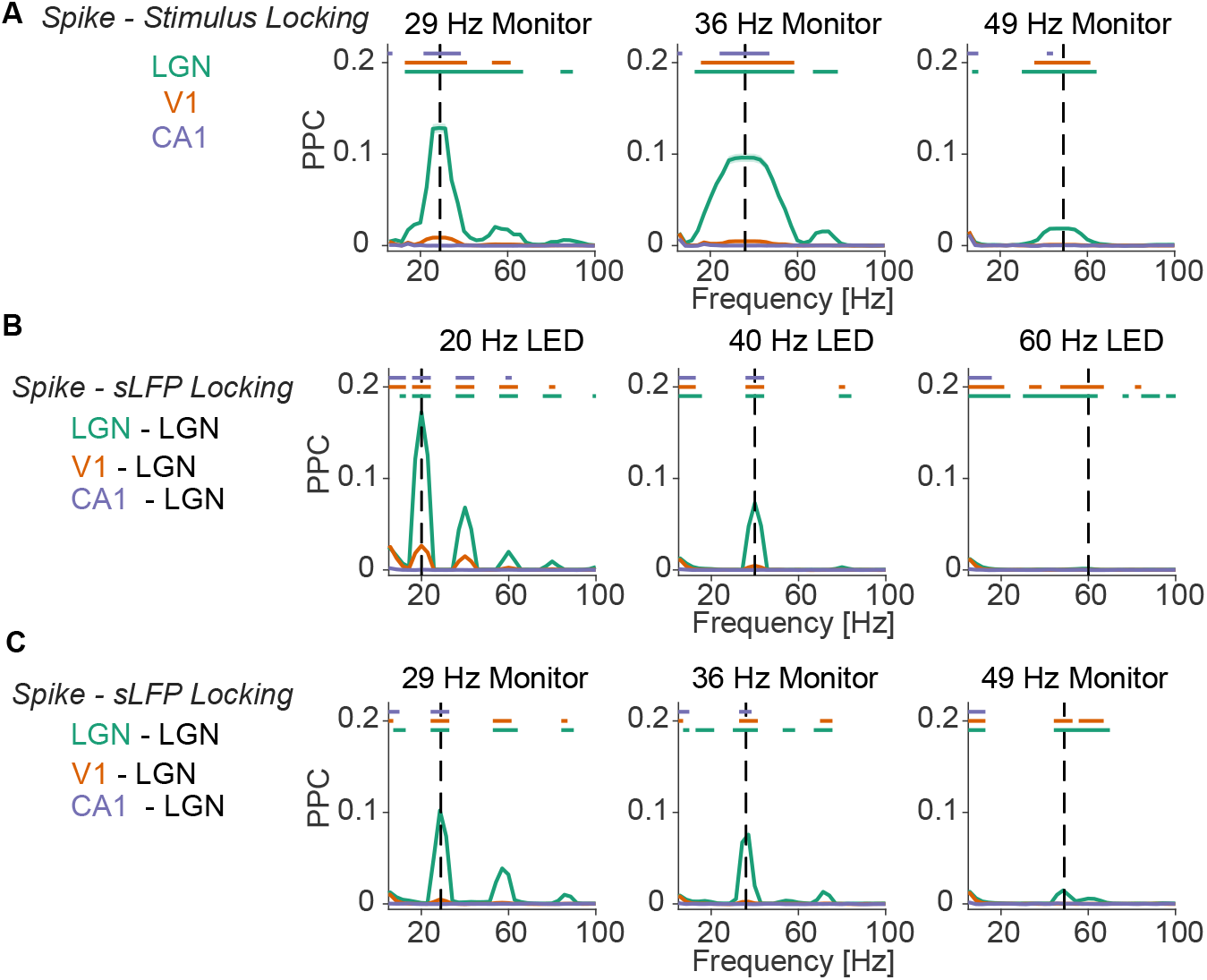
**(A)** Spike-field phase-locking during LED flicker stimulation (measured with PPC) for different combinations of spikes and sLFPs: Spikes in LGN (N=2386) to sLFPs in LGN (green); spikes in V1 (N=2091) to sLFP in LGN (orange); spikes in CA1 (N=636) to sLFP in LGN (blue). **(B)** Spike-field phaselocking during monitor flicker stimulation (measured with PPC) for different combinations of spikes and sLFPs: Spikes in LGN (N=1153 to sLFPs in LGN (green); spikes in V1 (N=815) to sLFP in LGN (orange); spikes in CA1 (N=125) to sLFP in LGN (blue). **(C)** Spike-stimulus phase-locking during monitor flicker stimulation (measured with PPC) of neurons in LGN (N=1153, green); V1 (N=815,orange) and CA1 (N=125, blue).

**Figure S4:**
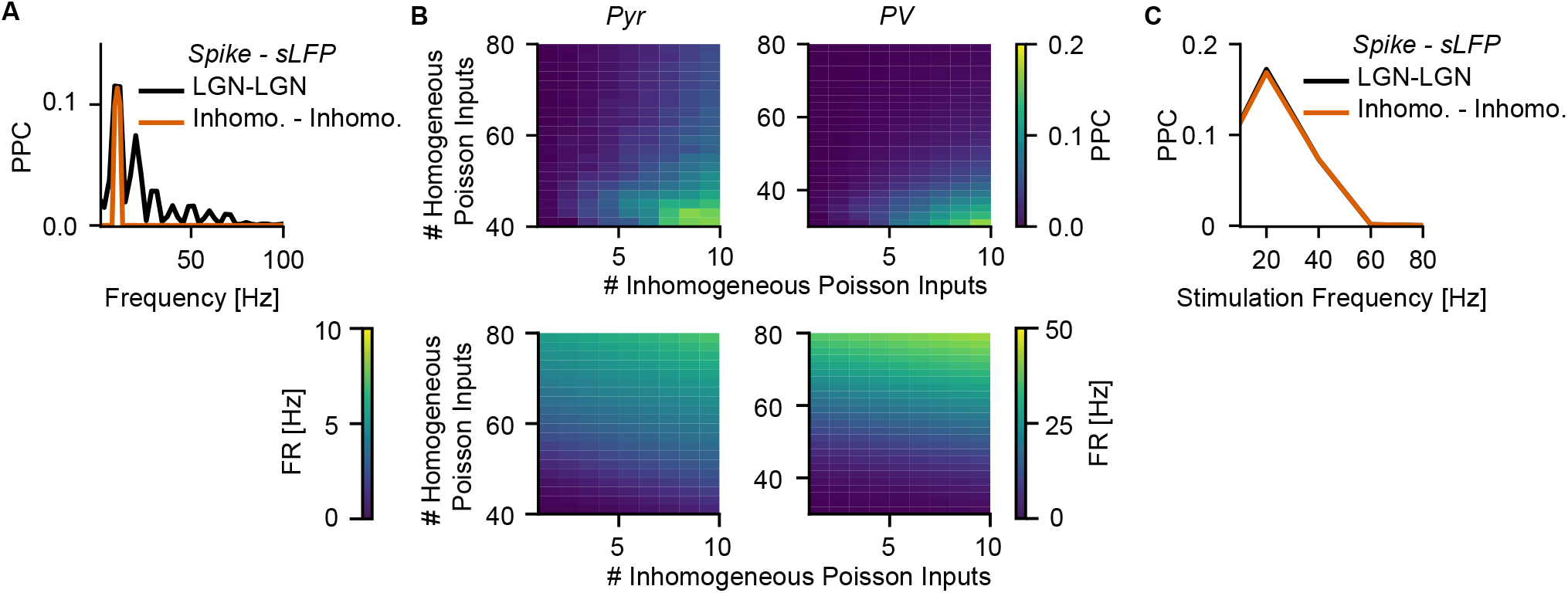
V1 multi-compartmental model during synaptic stimulation. **(A)** Spike -sLFP phase locking of LGN neurons during 10 Hz LED flicker stimulation to LGN population activity (black). Phase locking of inhomogeneous Poisson spiking input neurons (orange). Modulation strength adjusted to reproduce experimentally observed phase locking in LGN Inhomogeneous Poisson. **(B)** Grid scan of the number of homogeneous and inhomogeneous Poisson spiking input spike trains during 10 Hz stimulation. The top plots show phase locking (PPC) of Pyramidal (left) and PV+ (right) multi-compartmental model to inhomogeneous Poisson spiking input population. The bottom plots show the firing rates of corresponding neuron models. The number of homogeneous and inhomogeneous Poisson spiking input trains was optimized to fit the experimental results of BW and NW spiking neurons during 10 Hz LED stimulation. **(C)** Spike-sLFP phase locking (PPC) of single units in LGN to LGN population activity during stimulation at different frequencies (black). The modulation strength of inhomogeneous Poisson spiking input spike trains was adjusted to match experimental observations.

**Figure S5:**
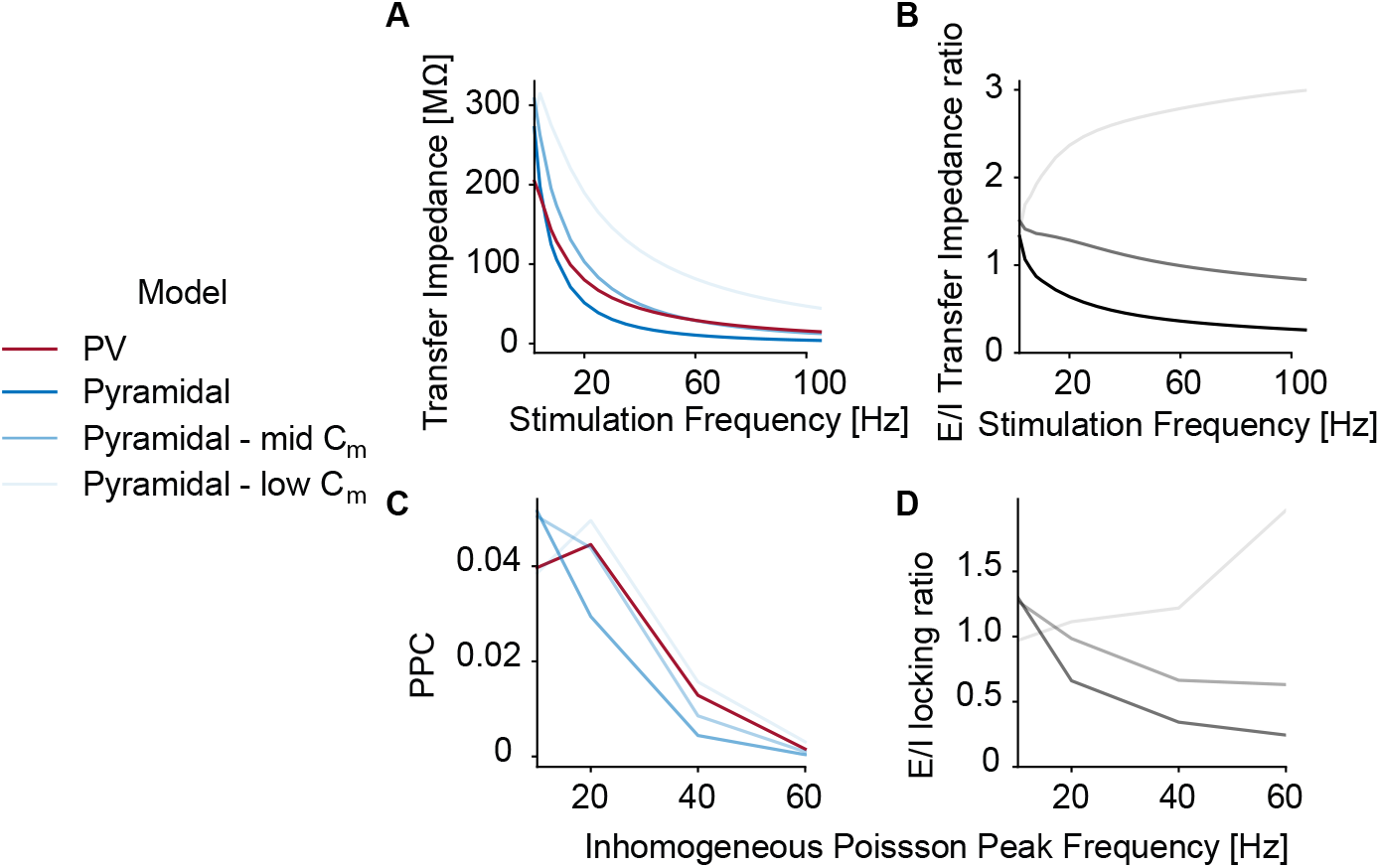
Effects of varying membrane capacitance on dendritic low-pass filtering. **(A)** Same simulations as in Figure 5A-D. Transfer impedance of PV+ and Pyramidal cell model during sinusoidal current stimulation between 1 and 100 Hz. Increasing the membrane capacitance in pyramidal cell dendrites resulted in a systematic increase in transfer impedance. **(B)** Ratio between the transfer impedance of the pyramidal cell model with scaled membrane capacitance and the PV+ neuron model. **(C)** Same simulations as in Figure 5H-K. Phase locking of PV+ and Pyramidal cell model with scaled membrane capacitance during synaptic stimulation with homogeneous and inhomogeneous spike trains. **(D)** Ratio between phase locking of the pyramidal cell model with scaled dendritic membrane capacitance and the PV+ cell model.

